# Large-scale brain correlates of sweet versus cocaine reward in rats

**DOI:** 10.1101/2022.06.01.494287

**Authors:** Magalie Lenoir, Sylvia Navailles, Youna Vandaele, Caroline Vouillac-Mendoza, Karine Guillem, Serge H. Ahmed

## Abstract

Cocaine induces many supranormal changes in neuronal activity in the brain, notably in learning- and reward-related regions, in comparison to nondrug rewards - a difference that is thought to contribute to its relatively high addictive potential. However, when facing a choice between cocaine and a nondrug reward (e.g., water sweetened with saccharin), most rats do not choose cocaine, as one would expect from the extent and magnitude of its global activation of the brain, but instead choose the nondrug option. We recently showed that cocaine, though larger in magnitude, is also an inherently more delayed reward than sweet water, thereby explaining why it has less value during choice and why rats opt for the more immediate nondrug option. Here we used a large-scale fos brain mapping approach to measure brain responses to each option in saccharin-preferring rats, with the hope to identify brain regions whose activity may explain the preference for the nondrug option. In total, fos expression was measured in 142 brain levels corresponding to 52 brain subregions and composing 5 brain macrosystems. Overall, our findings confirm in rats with a preference for saccharin that cocaine induces more global brain activation than the preferred nondrug option does. Only very few brain regions were uniquely activated by saccharin. They included regions involved in taste processing (i.e., anterior gustatory cortex) and also regions involved in processing reward delay and intertemporal choice (i.e., some components of the septohippocampal system and its connections with the lateral habenula).

## Introduction

Cocaine induces many supranormal changes in neuronal activity in the brain, notably in learning- and reward-related regions, in comparison to nondrug rewards (e.g., a palatable food) – a difference that is thought to contribute to its relatively high addictive potential and also to enduring brain neuroadaptations that may underlie chronic vulnerability to relapse. For instance, ample evidence in rats shows that intravenous cocaine causes dopamine surges in the ventral striatum that are much greater than those caused by a palatable food or drink (e.g., water sweetened with sucrose or saccharin) (reviewed in [1,2]). This difference not only persists but can even become larger with chronic drug use. Cocaine also causes more global neuronal activation of the brain as indicated by previous c-Fos brain mapping studies [3-5]. In one particularly relevant study in rats, self-administration of cocaine induced far greater changes in Fos expression than self-administration of sucrose did in all brain regions studied [3].

These established differences in brain neuronal activation between cocaine and nondrug rewards do not seem, however, to align well with other differences at the behavioral level, at least at first glance. Of particular importance, they do not predict rats’ choice between cocaine and nondrug rewards [6,7]. When facing such choice, most, and sometimes all rats, do not choose cocaine, as one would expect from the extent and magnitude of its global activation of the brain, but instead choose the nondrug option nearly exclusively and as a result stop using cocaine [1,8-22]. We recently found a possible resolution to this puzzle. Briefly, due to pharmacokinetics, rats consider cocaine during choice as a larger, albeit longer delayed, reward than sweet water and this despite its immediate intravenous delivery [2]. Once delivered, it takes tens of seconds for cocaine to produce its effects on ventral striatal dopamine. This inherently long delay would explain why rats attribute little value to cocaine in comparison to the more immediate alternative nondrug option during choice.

Here we used a large-scale fos brain mapping approach, as an indicator of neuronal activation [23,24], to try to identify in the brain some regions whose activity is associated with the most valued nondrug option, that is, water sweetened with saccharin in the present study. In total, we measured changes in fos expression in 142 brain levels corresponding to 52 different brain subregions and composing 5 brain macrosystems. Based on previous research, we expect that a large majority of brain regions will respond more to cocaine than to saccharin. However, we also expect to find some brain regions that will respond more or even uniquely to saccharin, notably brain regions known to be involved in processing and/or integrating reward magnitude and delay to compute value. One unique feature of the present study is that before measurement of brain region responses to each option, all rats had experience with self-administration of cocaine and saccharin, and they all preferred saccharin over cocaine during choice testing.

## Material and methods

### Subjects

A total of 58 young adult male Wistar rats (Charles River, L’Arbresle, France) were used (see Supplemental Information for more details about subjects, surgery, drugs and self-administration chambers).

### Initial operant training

Rats (n = 23) were initially trained on alternate days to press a lever (lever C) to receive cocaine intravenously (0.25 mg per injection) or to press a different lever (lever S) to obtain water-sweetened with 0.2% saccharin in an adjacent drinking cup (0.32 ml over a 20-s access). On average, they self-administered a total of 742±22 saccharin rewards and 422±16 cocaine rewards (see Experiment 1 in Supplemental Information for details about operant training).

### Discrete-trials choice procedure

After operant training, rats were tested under a discrete-trials choice procedure, as previously described [1,25]. Briefly, rats were allowed to choose during several consecutive daily sessions between lever C and lever S. Each daily choice session consisted of 12 discrete trials, spaced by 10 min, and divided into two successive phases, sampling (4 trials) and choice (8 trials). During sampling, each trial began with the presentation of one single lever in this alternate order: C – S – C – S. Lever C was presented first to prevent an eventual drug-induced taste aversion conditioning or negative affective contrast effects [1]. If rats responded on the available lever within 5 min, this triggered the immediate retraction of the lever, the immediate activation of the relevant syringe pump (i.e., saccharin or cocaine as described above) and the immediate illumination of the cue-light above the sampled lever for 40-s. If rats failed to respond within 5 min, the lever retracted until the next trial. Thus, during sampling, rats were allowed to evaluate each option separately before making their choice. Choice trials were identical to sampling trials, except that they began with the presentation of both levers S and C and ended with their simultaneous retraction. Rats had to respond on one of the two levers to make their choice and obtain the corresponding reward. During sampling and choice, the response requirement was set to 2 consecutive responses to avoid eventual accidental choice. A response on the alternate lever before satisfaction of the response requirement reset it. Response resetting occurred very rarely, however.

As previously reported [1,8,19,20], the large majority of rats (18 out of 22) preferred saccharin over cocaine (mean saccharin choices over the last 3 sessions: 91.4 ± 2.4%; range: 66.7-100.0%). Only few rats were indifferent (n = 2; saccharin choices: 54.2 and 45.8%) or preferred cocaine (n = 2; cocaine choices: 66.7 and 70.8%). Only the majority of saccharin-preferring rats were included in the large-scale Fos brain mapping experiment.

### Reward sampling test for brain activity analysis

Saccharin-preferring rats were distributed into 3 balanced groups (n = 5-7 per group) with similar average preference and prior history of cocaine and saccharin intakes (Table 1). During testing, one group was merely exposed to the operant cage for 90 min, with no programmed events (group CTL). This group controlled for any changes in brain activity due to the testing procedure and the exposure to the operant context. The two other groups were treated identically, except that they were given one single sampling trial for either saccharin (group SAC) or cocaine (group COC) as described above. After responding for either saccharin or cocaine, SAC and COC rats remained undisturbed in the operant context until the end of the 90-min period (see Experiment 1 in Supplemental information). Comparison of groups SAC and COC with group CTL should allow us to identify changes in brain activity specific to cocaine or saccharin reward (see below, for more information).

**Table 1:** Saccharin preferences during choice procedure and prior history of cocaine and saccharin intakes in the 3 experimental groups before reward sampling test for brain activity analysis.

| Group | Size | Prior SAC intake (ml/Kg BW) | Prior COC intake (mg/Kg BW) | SAC preference (%) |
| --- | --- | --- | --- | --- |
| CTL | n=6 | 530.0 ± 29.3 | 264.2 ± 23.4 | 93.8 ± 2.6 |
| SAC | n=5 | 565.4 ± 71.6 | 309.1 ± 18.9 | 96.7 ± 1.6 |
| COC | n=7 | 603.4 ± 41.6 | 263.5 ± 22.4 | 85.6 ± 5.1 |
|  |  | Factor: Experimental Group<br>F value: F(2,15)=0.65<br>P value: p=0.53 | Factor: Experimental Group<br>F value: F(2,15)=1.25<br>P value: p=0.32 | Factor: Experimental Group<br>F value: F(2,15)=2.25<br>P value: p=0.14 |

### c-Fos immunohistochemistry

Immunohistochemical procedures used in this study have been described in detail previously [26]. Following the final test session, we immediately anesthetized rats with sodium pentobarbital (120 mg/kg, i.p.) and perfused them intracardially with 150 ml of sodium phosphate buffer (PBS, 0.1M, pH 7.4), followed by 200 ml of 4% formaldehyde (VWR, France) in 0.1 M PBS. We perfused rats 90 minutes after the onset of reward delivery because previous research demonstrated that stimulated c-Fos protein peaked by 90-120 min [27]. We rapidly extracted brains after perfusion for subsequent single-label c-Fos immunohistochemistry (see Supplemental Information for detailed immunohistochemical procedure for Fos).

### Immunoreactivity counting

We already described in details how we generated a large-scale and high-resolution fos mapping of brain regions elsewhere [26]. For each rat, we counted c-Fos positive (Fos+) cells in both hemispheres from 2 to 6 sections (50 µm thick, 400 µm apart) covering the entire anteroposterior (AP) extent of each region of interest (ROI) within 5 major brain systems: 1) the cortico-thalamo-hippocampal system, 2) the striato-pallido-septal system, 3) the extended and basal amygdala system, 4) the hypothalamic-epithalamic-subthalamic system and 5) the midbrain, tegmentum and pons system (see the legend of the Table S1 in Supplemental Information for more details about the examined subregions in each major brain system). We collected sections either on 2 AP levels (one section per AP level, sections divided into anterior and posterior regions) or 3 AP levels (1 or 2 sections counted per AP level, identified as rostral (r), middle (m), and caudal (c)). We quantified c-Fos positive nuclei densities in 21 ROI composed of 52 subregions. For each rat, we averaged data from all sections per ROI and we calculated the mean ± standard error of the mean of these values (SEM) for each experimental group (COC, SAC and CTL) (see Supplemental Information for more details).

### Global measures of brain c-Fos expression

The c-Fos expression in the whole brain of each individual rat can be represented by the magnitude of an *n*-dimensional vector, with *n* corresponding to the total number of brain subregions studied (n = 52), and can be computed as follows:

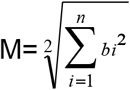

where *b*_*i*_ denotes the density of Fos+ cells in the *ith* brain region (i = 1 to *n*).

A similar approach was also used to measure the c-Fos expression in each of the 5 brain systems considered in the present study (see Section “Immunoreactivity counting” above). The only difference is that the dimensions of the vector representing each brain system corresponded to the specific subset of brain regions used to define it (n = 15 subregions for the cortico-thalamo-hippocampal, n = 15 for the striato-pallido-septal, n = 8 for the extended and basal amygdala, n = 7 for the hypothalamic-epithalamic-subthalamic and n = 7 for the midbrain, tegmentum and pons system).

### Statistical analysis

We determined sample sizes from previous studies [26]. Data are presented as mean values ± SEM. We performed statistical analyses by using Statistica, version 7.1 (Statsoft, Inc., Maisons-Alfort, France). All behavioral data were subjected to one-way analysis of variance (ANOVA) followed by appropriate *post-hoc* comparisons using the Tukey’s HSD test. Comparisons with a fixed theoretical level (e.g., indifference level at 50%) and between two means were conducted using one sample t-tests and unpaired t-tests, respectively. Numbers of Fos+ cells in brain regions were analyzed by using two- or three-way mixed ANOVA, followed by Tukey’s HSD *post-hoc* test. Comparisons between groups at each AP level were conducted using one-way ANOVA. Alpha level for detecting statistical significant differences was set to p < 0.05. Correction for multiple testing with a false discovery rate of 0.1 was applied using Benjamini-Hochberg procedure [28].

## Results

### Behavior during reward sampling test

During the reward sampling test, all rats from the SAC and COC groups responded on the available lever to obtain the corresponding reward within the 5-min allotted time. There was a trend that rats from group SAC responded faster for saccharin than rats from group COC for cocaine (3.5±0.7s versus 42.6±30.9s) but this difference did not reach statistical significance (t(10)=1.05, NS).

### Global brain changes of c-Fos expression

As expected, responding for cocaine, but not saccharin, induced a global increase in c-Fos expression in the brain (n = 52 subregions), as indicated by a significant increase in the magnitude of the brain vector above the control level (p=0.0018, post-hoc Tukey HSD, Fig. 1A). Though saccharin was the preferred reward, responding for it failed to induce such a global increase in c-Fos expression (p=0.11). A finer-grained anatomical analysis revealed that responding for cocaine induced c-Fos expression in 3 major brain systems: the cortico-thalamo-hippocampal system (p=0.014), the striato-pallido-septal system (p=0.017) and the hypothalamic-epithalamic-subthalamic system (p=0.0046) (see Fig. 1B, 1C and 1E). No significant change was observed in the other two brain systems considered, that is, the extended-basal amygdala system (Group: F(2,15)=2.04, NS) and the midbrain-tegmentum-pons system (p=0.13) (see Fig. 1D and 1F). In contrast, responding for saccharin only induced c-Fos expression in the cortico-thalamo-hippocampal system (p=0.023, Fig. 1B). No other significant changes were observed in the other brain systems.

**Figure 1:**
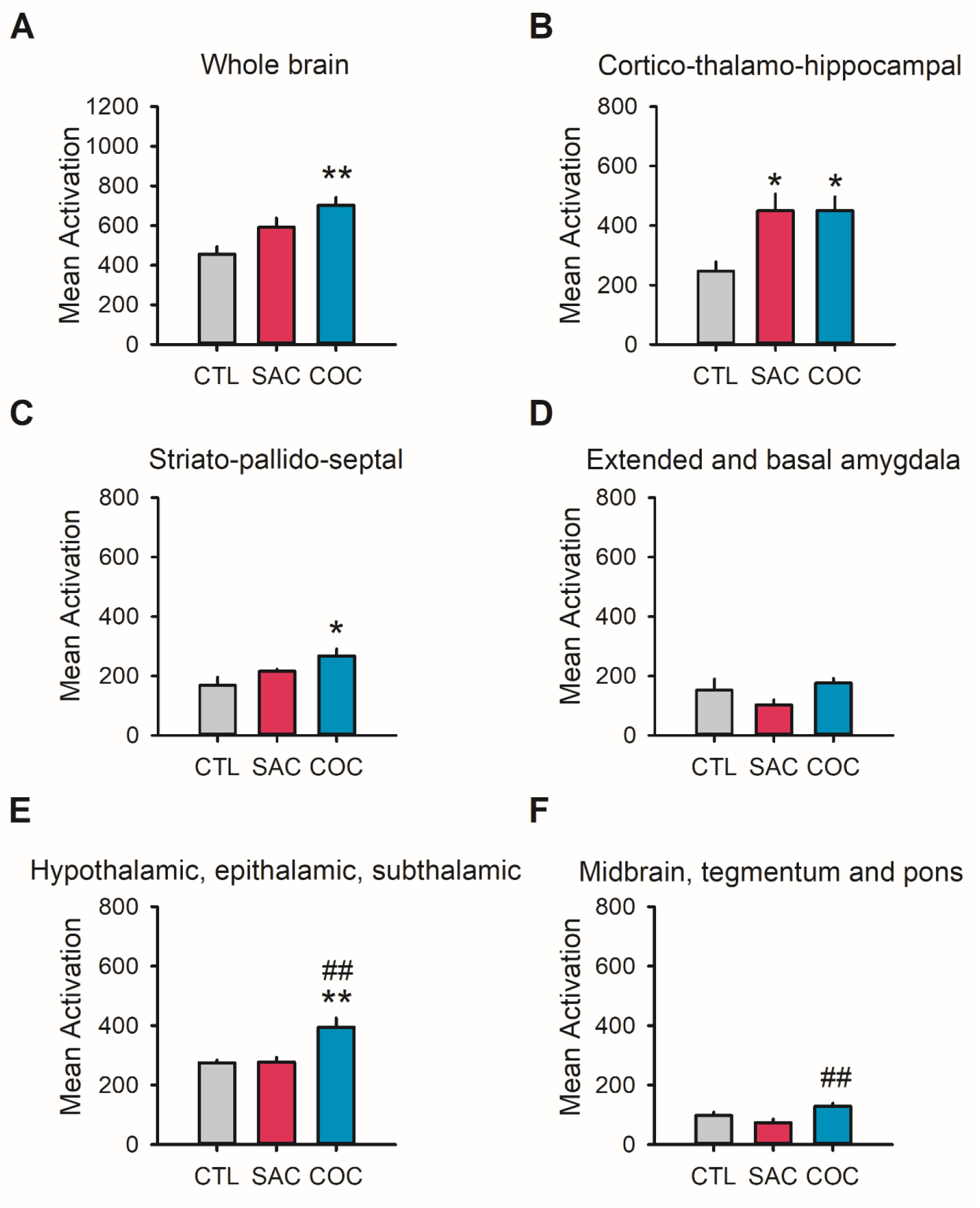
Whole-brain and regional brain changes of c-Fos expression induced by one sampling trial of cocaine or saccharin in saccharin-preferring rats previously trained to the choice procedure (Experiment 1). **A**. Bar graphs represent the magnitude of the multidimensional vector (± SEM) calculated from the individual densities c-Fos immunopositive nuclei counted within all the 52 subregions as a function of the experimental group (control (CTL, grey bars), saccharin (SAC, red bars) and cocaine (COC, blue bars), for more detail about calculation of the multidimensional vector, see Materials and Methods, section “Global measures of brain c-Fos expression”). **B, C, D, E, F**. Estimates of total c-Fos immunoreactivity by calculation of the magnitude of the multidimensional vector (± SEM) within the cortico-thalamo-hippocampal **(B)**, the striato-pallido-septal **(C)**, the extended and basal amygdala **(D)**, the hypothalamic, epithalamic, subthalamic **(E)** and the midbrain, tegmentum and pons systems **(F)**. * p<0.05, ** p<0.01, post-hoc Tukey’s HSD test, different from the CTL group. ## p<0.01, post-hoc Tukey’s HSD test, different from the SAC group. CTL: n=6, SAC: n=5, COC: n=7.

### Regional brain changes of c-Fos expression

With 3 experimental groups, there are a total of 12 possible patterns of group differences in regional brain c-Fos expression (Fig. S1), excluding the pattern of no group difference. Out of all these possible patterns, we actually observed only a subset of 6 patterns across all 52 brain regions studied (Fig. S1, bar graphs with grey backgrounds; see also Table S1 and Fig. S2 for representative photomicrographs of some of these patterns). Among the latter, the most frequent pattern by far corresponded to a reward-specific increase in c-Fos expression following responding for cocaine or saccharin (Table S1; Fig. S3). Of note, only few brain regions showed a nonspecific increase in c-Fos expression across rewards and will thus be ignored in the following analysis (Table S1; Fig. S4). A significant decrease in c-Fos expression was also rarely observed and when it was observed in a brain region, this was always following responding for saccharin and in association with an increase in c-Fos expression following responding for cocaine. In other words, the same brain region showed a pattern of opposite changes in c-Fos expression across drug and nondrug reward (see below, for more information and Table S1).

### Brain regions showing a saccharin reward-specific increase in c-Fos expression

As expected, we found a saccharin-specific increase in c-Fos expression in the anterior gustatory cortex (aGC) (aGC, SAC vs. CTL, p=0.0078, COC vs. CTL, NS, SAC vs. COC, p=0.0025; Fig. 2A). Such selective recruitment of the aGC is likely due to the sensory experience of the sweet taste of saccharin, confirming the sensitivity and selectivity of our brain mapping approach. Importantly, other non-sensory brain regions also exhibited a saccharin-specific increase in c-Fos expression. They included: the middle levels of the medial septum (mMS, SAC vs. CTL, p=0.047, COC vs. CTL, NS, SAC vs. COC, p=0.0066) and the vertical limb of the Diagonal Band of Broca (mvDBB, SAC vs. CTL, p=0.019, COC vs. CTL, NS, SAC vs. COC, p=0.021) (Fig. 2B). Interestingly, levels of c-Fos expression in these two brain regions were highly correlated following responding for saccharin (Pearson correlations, r=0.94, p=0.02) but not following responding for cocaine (r=-0.25, NS). In addition, increased c-Fos expression in the septal region was associated with an induction of c-Fos expression in the caudal part of the CA1 region of the dorsal hippocampus (cdCA1, SAC vs. CTL, p=0.020, Fig. 2B, insert). This finding suggests that responding for saccharin was associated with a selective activation of the septo-hippocampal system.

**Figure 2:**
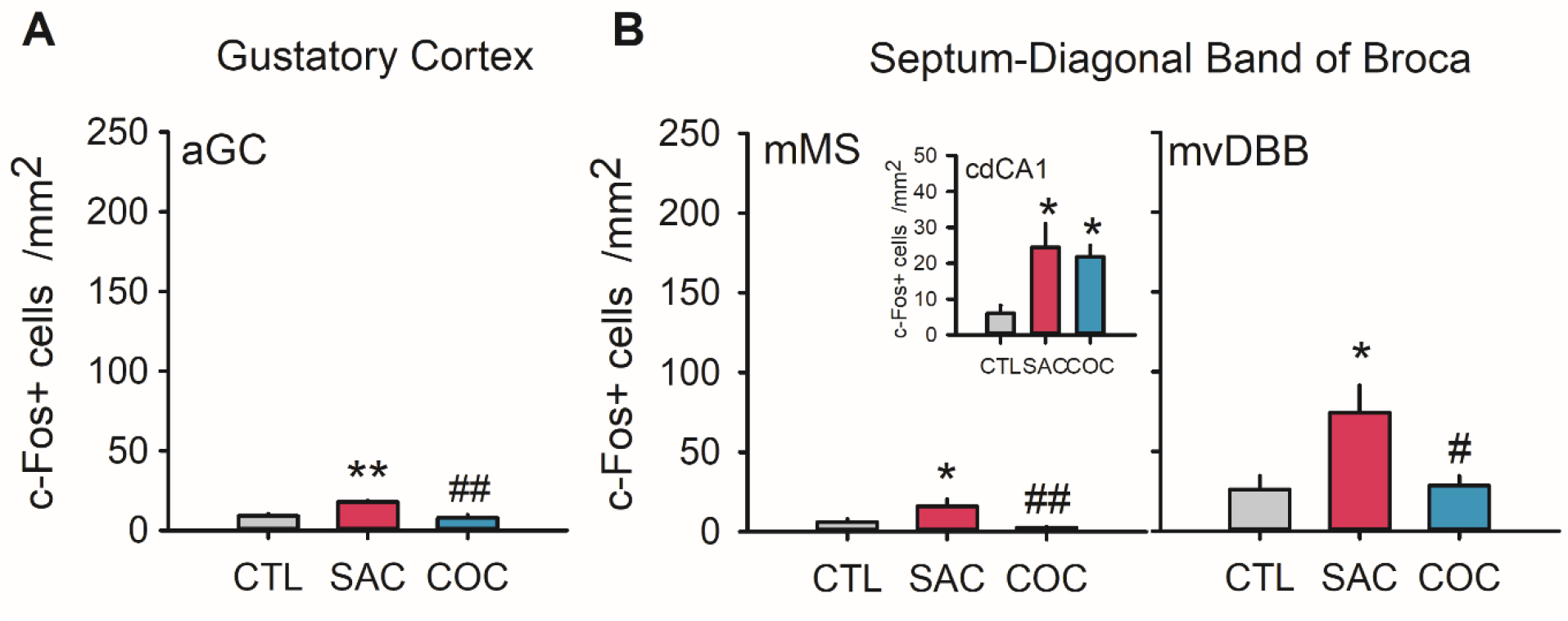
Saccharin-specific patterns of c-Fos activation observed in saccharin-preferring rats. **A, B**. Significant increases of c-Fos expression induced by a single trial of saccharin sampling in the anterior gustatory cortex **(A)** and in the medial septal-diagonal band of Broca (MS/vDBB) **(B)**. Bar graphs represent the mean (± SEM) density of Fos-positive cells (c-Fos+ cells /mm^2^) in rats exposed to a single trial of saccharin sampling (SAC group, red bars), a trial of cocaine sampling (COC group, blue bars) or to the operant cage only (CTL group, grey bars). **B, Insert**. Common activation of Fos expression at the caudal level of the dCA1 region by cocaine and saccharin samplings. * p<0.05, ** p<0.01, post-hoc Tukey’s HSD test, different from the CTL group. # p<0.05, ## p<0.01, post-hoc Tukey’s HSD test, different from the SAC group (CTL: n=6, SAC: n=5, COC: n=7). aGC, anterior level of the gustatory cortex; mMS, middle part of the medial septum; cdCA1, caudal level of the dorsal hippocampal CA1 region; mvDBB, middle part of the vertical limb of the Diagonal Band of Broca.

### Brain regions showing an opposite change in c-Fos expression across rewards

Only the middle level of the medial VTA (Group: F(2,15)=13.95, p=0.00038) and the rostral tVTA/RMTg (Group: F(2,15)=17.06, p=0.00014) (Fig. 3A), presented an opposite change in c-Fos expression following responding for cocaine and saccharin. Specifically, in both brain regions, c-Fos expression decreased following responding for saccharin while it increased following responding for cocaine (mMVTA, SAC vs. CTL, p=0.046, COC vs. CTL, p=0.044, SAC vs. COC, p=0.00041; tVTA/RMTg, SAC vs. CTL, p=0.033, COC vs. CTL, p=0.021, SAC vs. COC, p=0.00025). Of note, the rostral level of the lateral habenula (rLHb) also presented a saccharin reward-specific decrease in c-Fos expression (SAC vs. CTL, p=0.034, SAC vs. COC, p=0.020, Saccharin-specific Fos inhibition, Fig. 3B), but this was not associated with a significant cocaine-induced increase in c-Fos expression (COC vs. CTL, NS).

**Figure 3:**
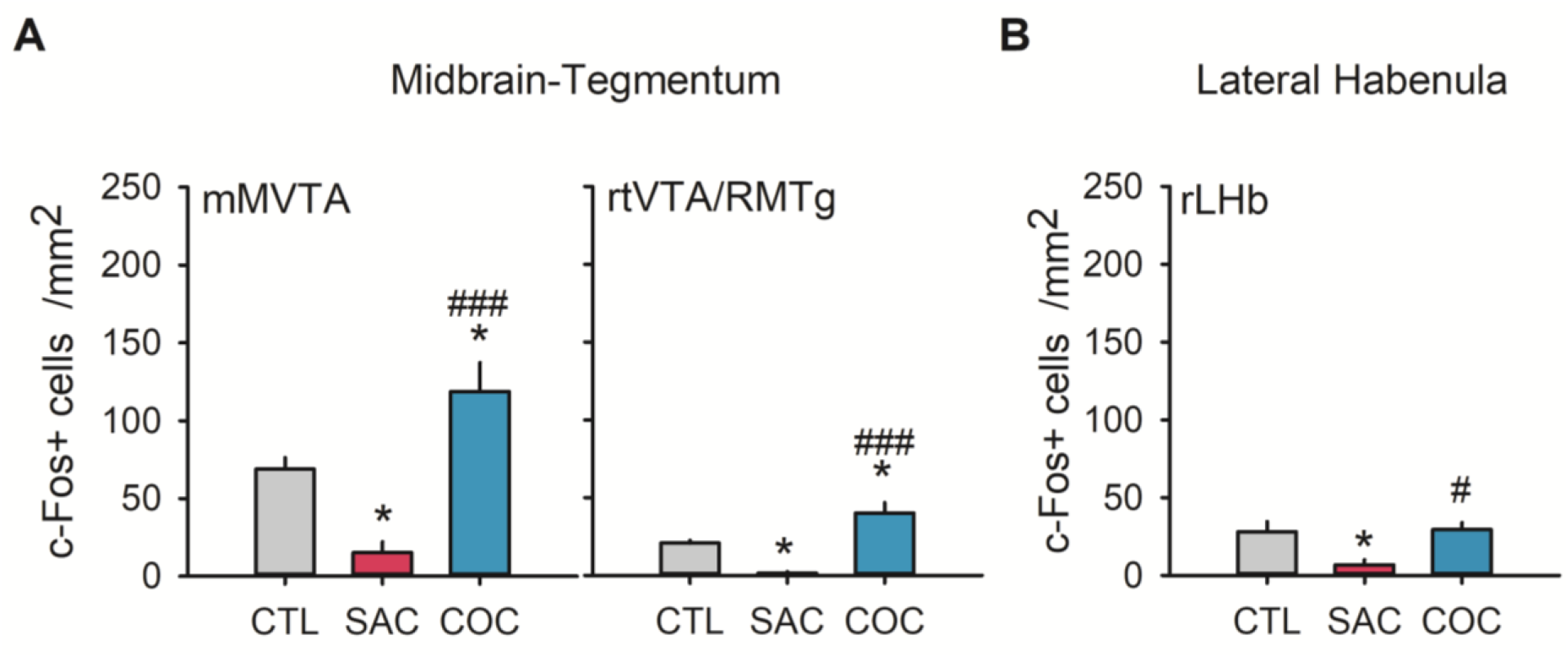
Opposite effects of saccharin and cocaine on c-Fos expression in midbrain nuclei of saccharin-preferring rats. **A**. Bar graphs represent mean (± SEM) density of Fos-positive cells (c-Fos+ cells /mm^2^) in restricted subregions of midbrain nuclei of rats exposed to a single trial of saccharin sampling (SAC group, n=5, red bars), a trial of cocaine sampling (COC group, n=7, blue bars) or only to the operant cage (CTL group, n=6, grey bars). Opposite effects of cocaine and saccharin on c-Fos expression are detected in the medial ventral tegmental area (VTA) and the tail of the VTA. **B**. A saccharin reward-specific decrease is also observed in the rostral level of the lateral habenula (rLHb), but this was not associated with significant cocaine-induced increase in c-Fos expression. * p<0.05, post-hoc Tukey’s HSD test, different from the CTL group. # p<0.05, ### p<0.001, post-hoc Tukey’s HSD test, different from the SAC group. mMVTA, middle level of the medial ventral tegmental area; rtVTA/RMTg, rostral part of the tail of the ventral tegmental area; rLHb, rostral part of the lateral habenula.

### c-Fos data generalizability

To better interpret some of the changes in c-Fos expression reported in the above experiment, we conducted a second experiment in a separate group of rats (n=28) that was similar to the first experiment but that also differed from it in several respects (see Experiment 2 in Supplemental Information for detailed method). First, rats were assigned to 3 independent groups, each with a different reward history. Briefly, all three groups were trained to respond under a discrete-trials sampling procedure to obtain either only cocaine (COC, n=12), only saccharin (SAC, n=8) or only the light cue that accompanied reward delivery in the first two groups (CTL, n=8). Second, by the end of training, the total number of rewards or light cue presentations obtained by each group was approximately equal before the final test for brain activity analysis (COC: 126 ± 15 rewards; SAC: 139 ± 9 rewards; CTL: 94 ± 5 light cues). Finally, during the final test, rats from each group were tested as during training, except that they were killed and perfused immediately at the end of the sampling session, that is, about 90 min after the first reward delivery, like in the first experiment. Thus, the second experiment was designed to measure reward-specific changes in c-Fos expression in rats with no prior experience with the two rewards and thus no experience of comparing their different values.

As observed in the first experiment, responding for cocaine, not for saccharin, induced a global increase in c-Fos expression in the brain (COC vs CTL: p=0.0034, Fig. 4A) and in several brain systems, including the cortico-thalamo-hippocampal system (p=0.0017), the striato-pallido-septal system (p=0.019), the midbrain-tegmentum-pons (p=0.00014, Fig. 4B, 4C and 4F) but not in the extended and basal amygdala system (Group: F(2,21)=0.96, NS; Fig. 4D) and the hypothalamic, epithalamic, subthalamic system (Group: F(2,21)=0.75, NS, Fig. 4E). In contrast, responding for saccharin only induced c-Fos expression in the striato-pallido-septal system, an effect that was comparable to that induce by cocaine (SAC vs CTL: p=0.024, Fig. 4C).

**Figure 4:**
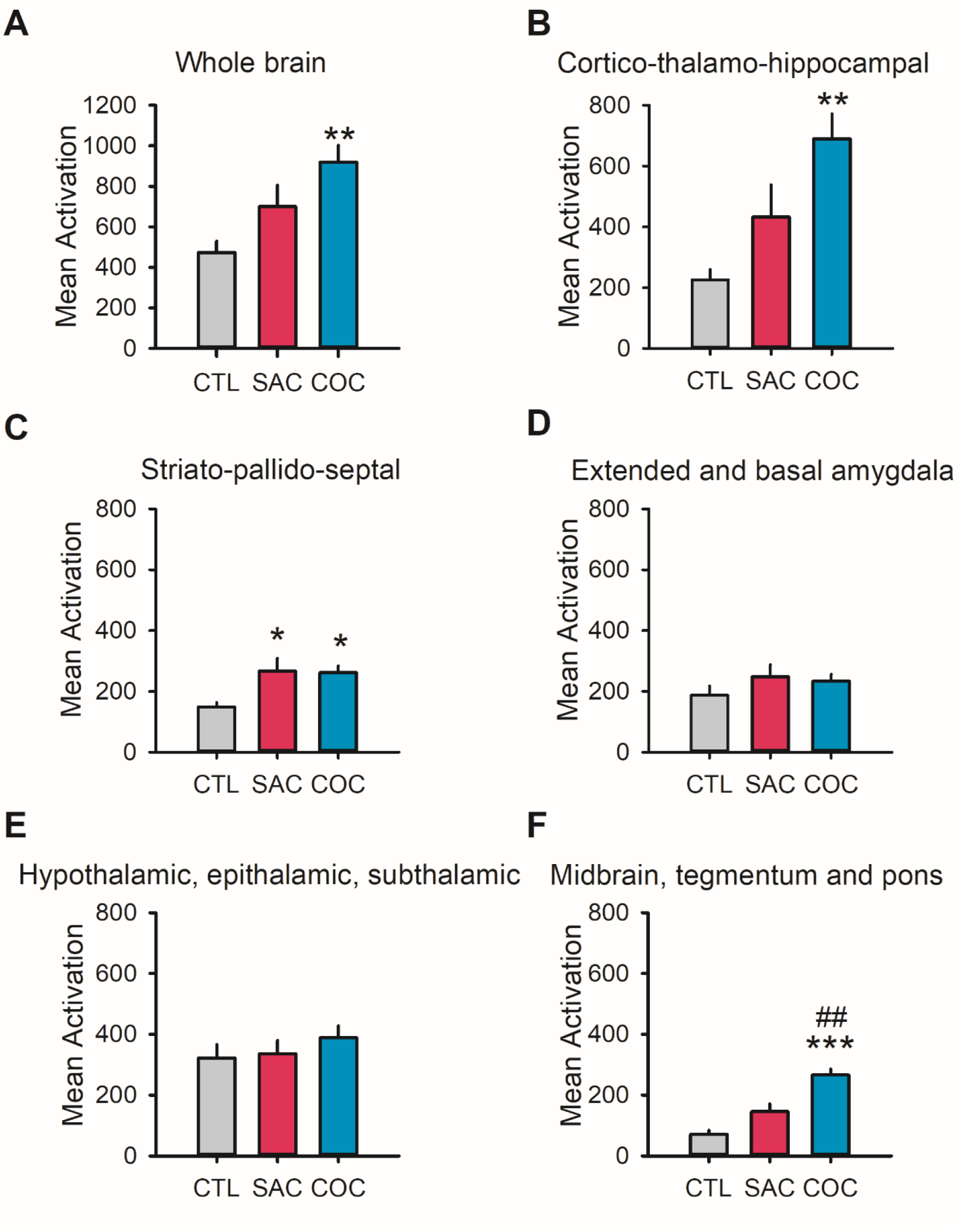
Whole-brain and regional brain changes of c-Fos expression induced by saccharin or cocaine in rats with previous experience with only one reward (Experiment 2). **A**. Bar graphs represent the magnitude of the multidimensional vector (± SEM) calculated from the individual densities c-Fos immunopositive nuclei counted within all the 52 subregions as a function of the experimental group (control (CTL, grey bars), saccharin (SAC, red bars) and cocaine (COC, blue bars), for more detail about calculation of the multidimensional vector, see Materials and Methods section). **B, C, D, E, F**. Estimates of total c-Fos immunoreactivity by calculation of the magnitude of the multidimensional vector (± SEM) within the cortico-thalamo-hippocampal **(B)**, the striato-pallido-septal **(C)**, the extended and basal amygdala **(D)**, the hypothalamic, epithalamic, subthalamic **(E)** and the midbrain, tegmentum and pons systems **(F)**. * p<0.05, ** p<0.01, *** p<0.001, post-hoc Tukey’s HSD test, different from the CTL group. ## p<0.01, post-hoc Tukey’s HSD test, different from the SAC group. CTL: n=7, SAC: n=7, COC: n=10.

In addition, as in the first experiment, a regional brain analysis revealed that responding for saccharin induced c-Fos expression into the septal nuclei (Fig. 5A, Septum-Diagonal Band of Broca, SAC vs. CTL: cMS, p=0.0061; cLS, p=0.0026) and in the caudal part of the vDBB (cvDBB, SAC vs. CTL, p=0.034, see the insert of Fig. 5A). We also noticed a saccharin-specific increase in c-Fos expression in the middle part of the ventromedial ventral pallidum (mvmVP, p=0.0054, Fig. 5B, Ventral Pallidum).

**Figure 5:**
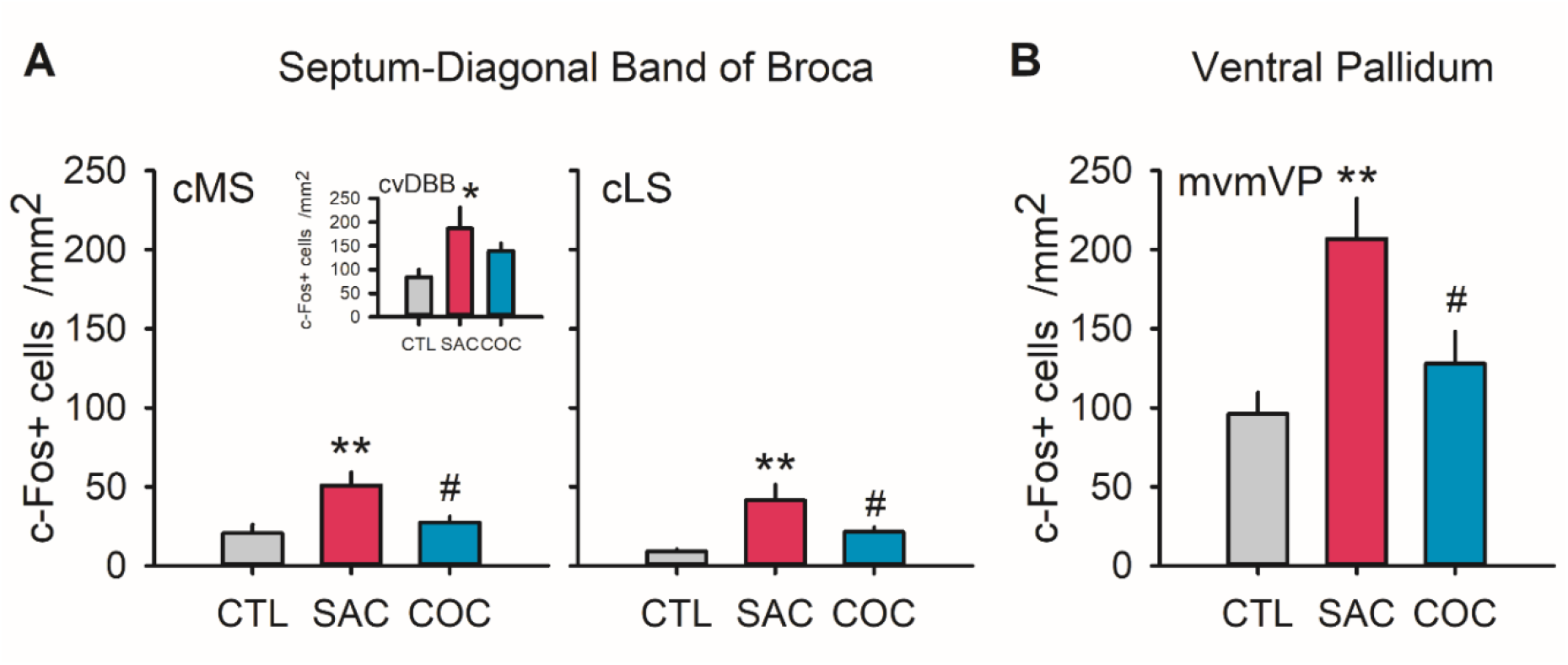
Saccharin-specific patterns of c-Fos activation in rats with previous experience with only one reward (Experiment 2). **A, B**. Bar graphs represent the mean (± SEM) density of Fos-positive cells (c-Fos+ cells /mm^2^) in rats exposed to trials of saccharin sampling (SAC group, red bars, n=7), cocaine sampling (COC group, blue bars, n=10) or only to the light cues that normally signal reward delivery (CTL group, grey bars, n=7). We observed significant increases of c-Fos expression induced by saccharin sampling into the medial septal-diagonal band of Broca complex (cMS/cvDBB and LS) **(A)** and the ventral pallidum **(B)**. * p<0.05, **p<0.01, post-hoc Tukey’s HSD test, different from the CTL group. # p<0.05, post-hoc Tukey’s HSD test, different from the SAC group. cMS, caudal level of medial septum; cvDBB, caudal level of the vertical limb of the Diagonal Band of Broca; cLS, caudal part of the lateral septum; mvmVP, middle level of the ventromedial ventral pallidum.

## Discussion

Our findings confirm and extend previous research [3-5]. Responding for cocaine, but not saccharin, induced a global increase in c-Fos expression in the brain, a difference seen in both rats with or without a prior experience of comparing and choosing between these two rewards. In particular, cocaine induced stronger c-Fos activations in the midbrain, tegmentum and pons system than those caused by saccharin. In addition, the rDMShell was gradually activated by the two rewards in saccharin-preferring rats, with stronger cocaine-induced c-Fos activation compared with that induced by saccharin (see Fig S4B). Responding for saccharin induced c-Fos expression in only one system, the cortico-thalamo-hippocampal system. Notably, saccharin activated equally or slightly more than cocaine the rostral level of the ventral (VO) and the middle level of the ventrolateral (VLO) orbitofrontal areas (Fig. S4A), a finding consistent with our previous research using in vivo recording of neuronal activity within the orbitofrontal cortex [19,29]. Not surprisingly, saccharin uniquely recruited the anterior gustatory cortex (aGC) which is involved in the sensory processing of sweet taste but also in other taste-related functions (see [30,31] for review). More interestingly, saccharin also uniquely recruited some components of the septohippocampal system that are not directly related to the sensory aspects of sweet taste. We will focus our discussion on this novel finding.

In both experiments, saccharin selectively activated the MS-vDBB subsystem of the septohippocampal system. MS-vDBB is a limbic structure involved in various cognitive and emotional processes, such as locomotion and motivational aspect of actions [32], arousal and attention [33], reward [34], feeding and consummatory behaviors [35,36], anxiety [37] and behavioral inhibition [38]. It should be pointed that MS and vDBB comprise at least 3 neuronal populations (cholinergic, GABAergic and glutamatergic neurons [39]) that are heterogeneously distributed on the rostrocaudal axis. It is unlikely that MS-vDBB activation in the present study involved anxiety because ventral, but not dorsal hippocampus is required for anxiety-related behaviors [32,40]. In addition, this saccharin-induced activation probably does not reflect the appetite suppressive role of medial septal glutamatergic neurons because downstream neurons from the paraventricular hypothalamus (a brain region that largely suppresses feeding and which is activated by MS glutamatergic neurons [35]) were not activated here. This saccharin-specific activation of MS-vDBB in both experiments further suggests that this structure is involved in processing saccharin reward. This is in accordance with the well-known role of MS in processing rewarding experiences and with the fact that rodents can work more strongly to obtain train electrical stimulations in the MS than in any other brain areas [34,41,42]. Interestingly, the MS encodes reward signals and positive motivational valence by sending GABAergic inhibitory projections to the LHb [43], a brain region also involved in choice between immediate and delayed rewards [44]. Consistent with this inhibitory control, we found that activation of MS-vDBB by saccharin was also associated with a specific decrease in c-Fos expression in the rLHb (Fig. 3B).

Another key role for the GABAergic MS-vDBB neurons has also been described in cognitive flexibility and perseverative inflexible-type behaviors [45-48]. It will be interesting to study if this system could be involved in the process reported previously in which preference for saccharin over cocaine becomes under habitual control after extended training in a drug choice setting [49,50]

Though MS-vDBB was activated by saccharin in both experiments, only one of its subcomponents (i.e., mMS-mvDBB-cdCA1) was activated in rats with a prior experience of choosing between saccharin and cocaine. A subpopulation of medial septal GABAergic neurons, expressing parvalbumin, displays spontaneous firing at theta frequency (4-12 Hz) in vivo [51]. This theta rhythmicity is then transmitted to the CA1 region of the hippocampus via septohippocampal projections [52] and to cortices, such as OFC [53]. The GABAergic septohippocampal pathway is involved in associative learning, in processing of the reward value and in modulating hippocampal rhythmic activities that are related to reward properties in mice [36]. This pathway is also implicated in integrating the delay that modulates the reward value [39]. Indeed, previous researches indicated that septal-lesioned rats were less likely to delay gratification, supporting a role of this pathway in mediating delayed reinforcement [54-56]. It has been recently shown that distinct subpopulations of CA1 hippocampal neurons encode delay and value information in mice performing a delay-discounting decision-making task [57]. In conclusion, this pathway is critically involved in delay discounting and intertemporal choice. Future research will need to elucidate the exact role of this pathway and the contribution of each neural population (GABA, glutamate and acetylcholine) in the decision-making process during choice between cocaine and saccharin.

This study has several limitations. First, animals experienced different behaviorally-relevant events during the final test, including exposure to the operant context, presentation and retraction of the reward-associated lever, lever pressing, illumination of the cue light associated with reward delivery and, finally, reward consumption and/or experience. Though we controlled for some of these events (e.g., context exposure and cue light), it is difficult to associate specifically changes in Fos expression in a brain region uniquely to one of these events to the exclusion of the others. However, the observation that responding for cocaine induces a global neuronal activation suggests that changes in Fos expression mainly reflects the acute experience of the reward itself. Second, though we found some brain regions that were uniquely activated by responding for saccharin, we probably missed other brain regions not detectable with postmortem c-Fos immunocytochemistry. This method has several well-known limitations [58-60]. For instance, because this technique has low temporal resolution, this can cause a loss of information about transient changes in neuronal activity. Moreover, c-Fos expression does not necessarily imply activation of neurons but can also be triggered by biochemical changes in intracellular signaling that are not necessarily associated with changes in neuronal activity [5,61]. Finally, c-Fos is less sensitive for marking cells, which are under net synaptic inhibition [58].

To conclude, our studies confirm previous research showing that cocaine induces more global brain activation than nondrug rewards and extend this finding to rats with a known preference for saccharin over cocaine. We also identified a unique association between some components of the septohippocampal system (i.e., MS-vDBB-dCA1) and responding for saccharin. Further research is needed to explore the role of these components during choice between cocaine and saccharin. Notably, it will be important to elucidate what are exactly the functions and causal role of the MS-vDBB-dCA1 in choice and preference between drug and nondrug rewards and whether this depends on a more general role in intertemporal choice.

## Acknowledgements

We thank Jean-Philippe Fougère and Eric Wattelet for administrative assistance. We also thank Sandra Dovero, Evelyne Doudnikoff, and Matthieu Bastide for technical advice, and Etienne Gontier from the Bordeaux Imaging Center (BIC) for giving us access to his laboratory facility for brain perfusion. Finally, we thank Dr Michel Engeln for his helpful comments on a previous version of the manuscript.

## Author contributions

SN and SHA conceived the project and designed the experiments. SN, YV, CVM and KG performed the experiments. ML, SN, YV and SHA analyzed the data. ML and SHA wrote the paper. All authors reviewed content and approved the final version of the manuscript.

## References

1 Lenoir M, Serre F, Cantin L, Ahmed SH. Intense sweetness surpasses cocaine reward. PloS one. 2007;2(8):e698.

2 Canchy L, Girardeau P, Durand A, Vouillac-Mendoza C, Ahmed SH. Pharmacokinetics trumps pharmacodynamics during cocaine choice: a reconciliation with the dopamine hypothesis of addiction. Neuropsychopharmacology : official publication of the American College of Neuropsychopharmacology. 2021;46(2):288–96.

3 Gao P, Limpens JH, Spijker S, Vanderschuren LJ, Voorn P. Stable immediate early gene expression patterns in medial prefrontal cortex and striatum after long-term cocaine self-administration. Addiction biology. 2017;22(2):354–68.

4 Park TH, Carr KD. Neuroanatomical patterns of fos-like immunoreactivity induced by a palatable meal and meal-paired environment in saline- and naltrexone-treated rats. Brain research. 1998;805(1-2):169–80.

5 Zahm DS, Becker ML, Freiman AJ, Strauch S, Degarmo B, Geisler S, et al. Fos after single and repeated self-administration of cocaine and saline in the rat: emphasis on the Basal forebrain and recalibration of expression. Neuropsychopharmacology : official publication of the American College of Neuropsychopharmacology. 2010;35(2):445–63.

6 Ahmed SH, Lenoir M, Guillem K. Neurobiology of addiction versus drug use driven by lack of choice. Current opinion in neurobiology. 2013;23(4):581–7.

7 Ahmed SH. Trying to make sense of rodents’ drug choice behavior. Progress in neuro-psychopharmacology & biological psychiatry. 2018;87(Pt A):3–10.

8 Cantin L, Lenoir M, Augier E, Vanhille N, Dubreucq S, Serre F, et al. Cocaine is low on the value ladder of rats: possible evidence for resilience to addiction. PloS one. 2010;5(7):e11592.

9 Augier E, Vouillac C, Ahmed SH. Diazepam promotes choice of abstinence in cocaine self-administering rats. Addiction biology. 2012;17(2):378–91.

10 Kerstetter KA, Ballis MA, Duffin-Lutgen S, Carr AE, Behrens AM, Kippin TE. Sex differences in selecting between food and cocaine reinforcement are mediated by estrogen. Neuropsychopharmacology : official publication of the American College of Neuropsychopharmacology. 2012;37(12):2605–14.

11 Perry AN, Westenbroek C, Becker JB. The development of a preference for cocaine over food identifies individual rats with addiction-like behaviors. PloS one. 2013;8(11):e79465.

12 Tunstall BJ, Kearns DN. Reinstatement in a cocaine versus food choice situation: reversal of preference between drug and non-drug rewards. Addiction biology. 2014;19(5):838–48.

13 Tunstall BJ, Riley AL, Kearns DN. Drug specificity in drug versus food choice in male rats. Experimental and clinical psychopharmacology. 2014;22(4):364–72.

14 Madsen HB, Ahmed SH. Drug versus sweet reward: greater attraction to and preference for sweet versus drug cues. Addiction biology. 2015;20(3):433–44.

15 Tunstall BJ, Kearns DN. Sign-tracking predicts increased choice of cocaine over food in rats. Behavioural brain research. 2015;281:222–8.

16 Tunstall BJ, Kearns DN. Cocaine can generate a stronger conditioned reinforcer than food despite being a weaker primary reinforcer. Addiction biology. 2016;21(2):282–93.

17 Vandaele Y, Cantin L, Serre F, Vouillac-Mendoza C, Ahmed SH. Choosing Under the Influence: A Drug-Specific Mechanism by Which the Setting Controls Drug Choices in Rats. Neuropsychopharmacology : official publication of the American College of Neuropsychopharmacology. 2016;41(2):646–57.

18 Kearns DN, Kim JS, Tunstall BJ, Silberberg A. Essential values of cocaine and non-drug alternatives predict the choice between them. Addiction biology. 2017;22(6):1501–14.

19 Guillem K, Ahmed SH. Preference for Cocaine is Represented in the Orbitofrontal Cortex by an Increased Proportion of Cocaine Use-Coding Neurons. Cerebral cortex. 2018;28(3):819–32.

20 Kim JS, Gunawan T, Tripoli CS, Silberberg A, Kearns DN. The effect of economy type on demand and preference for cocaine and saccharin in rats. Drug and alcohol dependence. 2018;192:150–57.

21 Bagley JR, Adams J, Bozadjian RV, Bubalo L, Ploense KL, Kippin TE. Estradiol increases choice of cocaine over food in male rats. Physiology & behavior. 2019;203:18–24.

22 Venniro M, Panlilio LV, Epstein DH, Shaham Y. The protective effect of operant social reward on cocaine self-administration, choice, and relapse is dependent on delay and effort for the social reward. Neuropsychopharmacology : official publication of the American College of Neuropsychopharmacology. 2021;46(13):2350–57.

23 Sagar SM, Sharp FR, Curran T. Expression of c-fos protein in brain: metabolic mapping at the cellular level. Science. 1988;240(4857):1328–31.

24 Morgan JI, Curran T. Stimulus-transcription coupling in neurons: role of cellular immediate-early genes. Trends in neurosciences. 1989;12(11):459–62.

25 Lenoir M, Augier E, Vouillac C, Ahmed SH. A choice-based screening method for compulsive drug users in rats. Current protocols in neuroscience. 2013;Chapter 9:Unit 9 44.

26 Navailles S, Guillem K, Vouillac-Mendoza C, Ahmed SH. Coordinated Recruitment of Cortical-Subcortical Circuits and Ascending Dopamine and Serotonin Neurons During Inhibitory Control of Cocaine Seeking in Rats. Cerebral cortex. 2015;25(9):3167–81.

27 Chaudhuri A, Zangenehpour S, Rahbar-Dehgan F, Ye F. Molecular maps of neural activity and quiescence. Acta neurobiologiae experimentalis. 2000;60(3):403–10.

28 Benjamini Y, Hochberg Y. Controlling the False Discovery Rate: A Practical and Powerful Approach to Multiple Testing. Journal of the Royal Statistical Society. 1995;57(No. 1):289–300.

29 Guillem K, Ahmed SH. Reorganization of theta phase-locking in the orbitofrontal cortex drives cocaine choice under the influence. Scientific reports. 2020;10(1):8041.

30 McCaughey SA. The taste of sugars. Neuroscience and biobehavioral reviews. 2008;32(5):1024–43.

31 Vincis R, Fontanini A. Central taste anatomy and physiology. Handbook of clinical neurology. 2019;164:187–204.

32 Mocellin P, Mikulovic S. The Role of the Medial Septum-Associated Networks in Controlling Locomotion and Motivation to Move. Frontiers in neural circuits. 2021;15:699798.

33 Wu M, Zhang Z, Leranth C, Xu C, van den Pol AN, Alreja M. Hypocretin increases impulse flow in the septohippocampal GABAergic pathway: implications for arousal via a mechanism of hippocampal disinhibition. The Journal of neuroscience : the official journal of the Society for Neuroscience. 2002;22(17):7754–65.

34 Olds J, Milner P. Positive reinforcement produced by electrical stimulation of septal area and other regions of rat brain. J Comp Physiol Psychol. 1954;47(6):419–27.

35 Sweeney P, Li C, Yang Y. Appetite suppressive role of medial septal glutamatergic neurons. Proceedings of the National Academy of Sciences of the United States of America. 2017;114(52):13816–21.

36 Vega-Flores G, Gruart A, Delgado-Garcia JM. Involvement of the GABAergic septo-hippocampal pathway in brain stimulation reward. PloS one. 2014;9(11):e113787.

37 Zhang Y, Jiang YY, Shao S, Zhang C, Liu FY, Wan Y, et al. Inhibiting medial septal cholinergic neurons with DREADD alleviated anxiety-like behaviors in mice. Neuroscience letters. 2017;638:139–44.

38 McNaughton N, Gray JA. Anxiolytic action on the behavioural inhibition system implies multiple types of arousal contribute to anxiety. Journal of affective disorders. 2000;61(3):161–76.

39 Robinson JC, Brandon MP. Skipping ahead: A circuit for representing the past, present, and future. eLife. 2021;10.

40 Bannerman DM, Rawlins JN, McHugh SB, Deacon RM, Yee BK, Bast T, et al. Regional dissociations within the hippocampus--memory and anxiety. Neuroscience and biobehavioral reviews. 2004;28(3):273–83.

41 Ball GG, Gray JA. Septal self-stimulation and hippocampal activity. Physiology & behavior. 1971;6(5):547–9.

42 Cazala P, Galey D, Durkin T. Electrical self-stimulation in the medial and lateral septum as compared to the lateral hypothalamus: differential intervention of reward and learning processes? Physiology & behavior. 1988;44(1):53–9.

43 Shen L, Zhang GW, Tao C, Seo MB, Zhang NK, Huang JJ, et al. A bottom-up reward pathway mediated by somatostatin neurons in the medial septum complex underlying appetitive learning. Nature communications. 2022;13(1):1194.

44 Stopper CM, Floresco SB. What’s better for me? Fundamental role for lateral habenula in promoting subjective decision biases. Nature neuroscience. 2014;17(1):33–5.

45 Bortz DM, Gazo KL, Grace AA. The medial septum enhances reversal learning via opposing actions on ventral tegmental area and substantia nigra dopamine neurons. Neuropsychopharmacology : official publication of the American College of Neuropsychopharmacology. 2019;44(13):2186–94.

46 Dwyer TA, Servatius RJ, Pang KC. Noncholinergic lesions of the medial septum impair sequential learning of different spatial locations. The Journal of neuroscience : the official journal of the Society for Neuroscience. 2007;27(2):299–303.

47 Pang KC, Jiao X, Sinha S, Beck KD, Servatius RJ. Damage of GABAergic neurons in the medial septum impairs spatial working memory and extinction of active avoidance: effects on proactive interference. Hippocampus. 2011;21(8):835–46.

48 Feldon J, Gray JA. Effects of medial and lateral septal lesions on the partial reinforcement extinction effect at short inter-trial intervals. The Quarterly journal of experimental psychology. 1979;31(Pt 4):675–90.

49 Vandaele Y, Guillem K, Ahmed SH. Habitual Preference for the Nondrug Reward in a Drug Choice Setting. Frontiers in behavioral neuroscience. 2020;14:78.

50 Vandaele Y, Lenoir M, Vouillac-Mendoza C, Guillem K, Ahmed SH. Probing the decision-making mechanisms underlying choice between drug and nondrug rewards in rats. eLife. 2021;10.

51 Tsanov M. Differential and complementary roles of medial and lateral septum in the orchestration of limbic oscillations and signal integration. The European journal of neuroscience. 2018;48(8):2783–94.

52 Buzsaki G. Theta oscillations in the hippocampus. Neuron. 2002;33(3):325–40.

53 Knudsen EB, Wallis JD. Closed-Loop Theta Stimulation in the Orbitofrontal Cortex Prevents Reward-Based Learning. Neuron. 2020;106(3):537–47 e4.

54 Gorenstein EE, Newman JP. Disinhibitory psychopathology: a new perspective and a model for research. Psychological review. 1980;87(3):301–15.

55 White NM. Effects of septal lesions on responding for delayed brain stimulation reinforcement. Brain research. 1974;65(2):185–93.

56 Rawlins JN, Feldon J, Butt S. The effects of delaying reward on choice preference in rats with hippocampal or selective septal lesions. Behavioural brain research. 1985;15(3):191–203.

57 Masuda A, Sano C, Zhang Q, Goto H, McHugh TJ, Fujisawa S, et al. The hippocampus encodes delay and value information during delay-discounting decision making. eLife. 2020;9.

58 Kovacs KJ. Measurement of immediate-early gene activation-c-fos and beyond. Journal of neuroendocrinology. 2008;20(6):665–72.

59 Kovacs KJ. c-Fos as a transcription factor: a stressful (re)view from a functional map. Neurochemistry international. 1998;33(4):287–97.

60 McReynolds JR, Christianson JP, Blacktop JM, Mantsch JR. What does the Fos say? Using Fos-based approaches to understand the contribution of stress to substance use disorders. Neurobiology of stress. 2018;9:271–85.

61 Harlan RE, Garcia MM. Drugs of abuse and immediate-early genes in the forebrain. Molecular neurobiology. 1998;16(3):221–67.

