## Supplemental information for "Large-scale brain correlates of sweet versus cocaine reward in rats"

### Supplementary information

#### Supplementary Methods:

##### Subjects and Housing

Young adult male Wistar rats (Charles River, L'Arbresle, France), weighing 200-225 g upon arrival, were used. Rats were housed in groups of two per cage and maintained in a light- (10h-14h reverse light-dark cycle) and temperature-controlled vivarium (21 ± 2°C). All behavioral testing occurred during the dark phase of the light-dark cycle. Standard rat chow A04 (SAFE, Scientific Animal Food and Engineering, Augy, France, see [1] for nutrients composition) and tap water were freely available in the home cages throughout the duration of the experiment (i.e., rats were neither food- nor water-restricted). No synthetic or refined sugar was added. Home cages were enriched with a nylon gnawing bone and a cardboard tunnel (Plexx BV, The Netherlands). All experiments were carried out in accordance with institutional and international standards of care and use of laboratory animals [UK Animals (Scientific Procedures) Act, 1986; and associated guidelines; the European Communities Council Directive (2010/63/UE, 22 September 2010) and the French Directives concerning the use of laboratory animals (decree 2013-118, 1 February 2013)]. All experiments have been approved by the Committee of the Veterinary Services Gironde, agreement number A33-063-922. We excluded in total 12 rats (Experiment 1: n=8, Experiment 2: n=4) due to catheter failure (Experiment 1: n=5), death during intravenous surgery (Experiment 1: n=2), software issue during the test day for subsequent c-Fos immunohistochemical analysis (Experiment 1: n=1), failure to respond for cocaine during the final test (Experiment 2: n=1) and to poor brain perfusion (Experiment 2: n=3), thereby leaving 46 rats for final analysis.

##### Surgery

Rats were anesthetized with a mixture of ketamine (110 mg/kg, i.p., Bayer Pharma, Lyon, France) and xylazine (15 mg/kg, i.p., Merial, Lyon, France). They were then implanted with indwelling silastic catheters (Dow Corning Corporation, Michigan, USA) into the right jugular vein that exited the skin in the middle of the back about 2 cm below the scapulae under aseptic conditions. After surgery, we flushed catheters daily with 0.2 ml of a sterile antibiotic solution containing heparinized saline (280 IU / ml) and ampicilline (Panpharma, Fougères, France) to maintain patency. Behavioral testing commenced at least 8 days after surgery. When a catheter leakage was suspected, we checked its patency by an intravenous administration of Etomidate (1 mg/kg, Braun Medical, Boulogne-Billancourt, France), a short-acting non-barbiturate anesthetic.

#### Drugs

Cocaine hydrochloride (Coopération Pharmaceutique Française, Melun, France) was dissolved in a solution of NaCl (0.9%) which was filtered through a syringe filter (0.22 µm) and kept at room temperature (RT) in a 500-ml sterile bag. Drug doses were expressed as the weight of the salt. We dissolved sodium saccharin (Sigma-Aldrich, St Quentin-Fallavier, France) in tap water at RT. Sweet solutions were renewed each day.

#### Self-administration chambers

Rats were trained in fourteen identical operant chambers (30×40×36 cm) for all behavioral testing (Imétronic, Pessac, France). All chambers were individually enclosed in wooden cubicles equipped with a white noise speaker (45±6 dB) for sound-attenuation and an exhaust fan for ventilation. The apparatus has been extensively described elsewhere [2]. Briefly, each chamber was equipped with two retractable metal levers on opposite panels of the chamber, and a corresponding white cue light was positioned above each lever. A drinking receptacle or cup was located 6.5 cm from the left of each retractable lever on the same panel and 6 cm above the grid floor. A lickometer circuit allowed for monitoring and recording of receptacle contacts. One syringe pump delivered saccharin solution to the drinking receptacle via Silastic tubing (Dow Corning Corporation, MI, USA). The other syringe pump delivered drug solution through Tygon tubing (Cole Parmer,Vernon Hills, IL, USA) connected via a single channel liquid swivel (Lomir Biomedical Inc., Quebec, Canada) to a cannula connector (Plastics One, Roanoke, VA, USA) on the back of the animal.

#### Experiment 1

##### Initial cocaine and saccharin self-administration training

Rats were first habituated to the operant chambers during two consecutive 2h30 daily sessions. During habituation, all levers were retracted and there was no programmed reward contingency. We then trained rats to press the left lever (lever S) for saccharin solution and the right lever (lever C) for intravenous cocaine injection on alternate days under a fixed-ratio 1 time-out 20-s schedule of reinforcement during 16 daily sessions. We chose a unit dose of 0.25 mg/infusion delivered over 4 s for intravenous cocaine self-administration training based on our previous studies [3-5]. A single press on the lever S was rewarded by a 20-s access to a drinking spout that delivered discrete volumes (0.04 ml) of a solution of sodium saccharin at a near optimal concentration of 0.2% [6-8]. Saccharin delivery was signaled by illumination of the 20s cue-light above the corresponding lever. The first 2 volumes were delivered freely during the first 3 s to fill the drinking cup; subsequent volumes were obtained by licking (1 volume per 20 licks in about 2.8 s for the remaining 17s). Cocaine and saccharin self-administration sessions ended after rats had earned a maximum of 20 rewards or after 2h and 2h30 had elapsed for saccharin and cocaine sessions, respectively. The beginning of each session started with the first lever press or after 10 min elapsed, whatever the reward tested. We conducted a minimum of 5 daily training sessions per reinforcer and we continued the training until each individual rat earned exactly 100 rewards for each type of reward. Following this initial training, they self-administered cocaine and saccharin during 34 additional daily sessions under different experimental conditions for an unrelated research question before being submitted to the mutually exclusive discrete choice procedure.

##### Reward sampling test for brain activity analysis

On the final test day, we exposed saccharin-preferring rats to one single sampling trial of cocaine (COC group, n=7), or to one sampling trial of saccharin (SAC group, n=5), whereas rats from the control group (CTL group, n=6) were only exposed to the operant cage. Following 10 min after the beginning of the test, we inserted the right lever in the COC group or the left lever in the SAC group. The lever remained extended for 5 minutes or until rats pressed the lever twice. In the CTL group, the both levers were retracted, no reward and no cue-light above the levers were delivered. This last group controlled for eventual neuronal correlates of the contextual stimuli. If rats of the COC or SAC groups responded on the lever within 5 minutes, they received the reward corresponding with the selected lever. Reward delivery was signaled by the retraction of the lever and a 40-s illumination of the cue-light above the selected lever and the retraction of the lever. If the rat failed to respond on the lever, the lever retracted with no reward delivery. It is noteworthy that all rats pressed the lever within the allocated time of 5 min. The final test ended 90 minutes after rats received the corresponding reward or placed in the context.

#### Experiment 2

##### Initial cocaine or saccharin self-administration training

In this experiment, rats were first habituated to the operant chambers during two consecutive 3h daily sessions. Rats were then randomly allocated into 3 different groups according to the type of reward received. The first group of rats was only allowed to self-administer cocaine (COC group, n=12, 0.25 mg, i.v., left lever) under a FR1 TO20s reinforcement schedule during 5 consecutive sessions. The second group had access to saccharin reward (SAC group, n=8, 20-s of access to water containing 0.2% saccharin, left lever) under the same schedule of reinforcement during the same 5 days. Finally, the last control group was only exposed to the operant cage (i.e., self-administration chamber, left lever extended, white noise, each lever press resulted in illumination of the 20s cue-light above the lever; CTL group, n=8). All the other experimental conditions are similar to those applied in self-administration training of the Experiment 1. Rats were limited to obtaining 20 reinforcers per session and/or each session ended after 3 h. After completing the training phase, we conducted the discrete-trials sampling procedure on the subsequent day.

##### Discrete-trials sampling procedure

We used a sampling procedure, in which the total number of trials per session was not set and depended on the rat's performance, during 9 daily sessions. Because the duration of sessions was set to 100 min and discrete trials were spaced by 10 min, each session consisted of a minimum of 6 discrete trials for a rat which did not complete any of its trials, while a maximum of 9 discrete trials could be done by a rat which completed 100% of sampling trials. In this sampling procedure, following 10 min after the beginning of the test, each trial began with the presentation of the left lever and rats had to press it within 5 min to obtain the corresponding reward (i.e, an intravenous injection of 0.25 mg cocaine over 4s for the COC group, a 20-s access to 0.2% saccharin for the SAC group or an illumination of the 40s cue-light above the lever for the CTL group). If rats responded on the lever, reward delivery was signaled by the retraction of the lever and a 40-s illumination of the cue-light above the lever. A failure to respond on the lever within 5 min resulted in its retraction, as well as no cue-light appeared and no reward was delivered in COC and SAC groups. However, no response on the lever within 5 min from the CTL group caused the retraction of the lever with the concomitant 40-s illumination of the cue-light above the lever. CTL group was exposed to 6-7 cue-light presentations over the 100-min session, thus having approximately the same exposure to discrete and contextual stimuli than rats from COC and SAC groups. This CTL group served as a direct control for eventual neuronal activation by the light presentation alone and lever pressing per se.

##### Reward sampling test for brain activity analysis

During the final test, rats were tested on a discrete trial 100-min sampling session similar to the 9 training discrete-trials sessions. Sessions consisted of 6-9 discrete trials spaced by 10 min, and ended after 100 min had elapsed. Rats were anesthetized and perfused immediately at the end of the 100-min session so that the time of sacrifice corresponded to 90 min after the first lever presentation/reward delivery. Only one rat failed to respond for cocaine during the test and was excluded from the experiment.

#### c-Fos immunohistochemistry

Collected brains were post-fixed in the same fixative solution at 4°C, then cryoprotected at 4 °C in 20% sucrose solution (Sigma-Aldrich, St. Quentin Fallavier, France) for at least 3 days. We then immersed brains in SnapFrost® system (Alphelys, France) containing cold isopentane at -55°C. Frozen brains were stored at -80°C until sectioning. Freezing 50 µm-tick coronal sections were cut on a cryostat (Leica CM 3050S) at -20°C and stored at 4°C in 0.1M PBS containing 0.03 % (w/v) of sodium azide. After being washed with PBS (0.1M, 3 times for 10 min) and incubated with $H_2O_2$ 0.3% for 30 min, free-floating sections were treated with blocking solution (3% normal goat serum in 0.1 M PBS containing 0.3% Triton X-100) for 1h at RT. The sections were then incubated with the primary rabbit polyclonal anti-c-Fos antibody for 48 h at 4 °C (1:8000; sc-52, Santa-Cruz Biotechnology Santa Cruz, CA), washed in PBS and incubated for 2h at RT with biotinylated goat anti-rabbit IgG (BA-1000: 1:200; Vector Laboratories, distributed by Clinisciences, France). Tissue sections were further processed using avidin–biotin–peroxidase complex (ABC, 1:200 for 2h at RT; ABC Vectastain Elite Kit, PK-6100; Vector laboratories) and 3,3′-diaminobenzidine detection (DAB: 0.05% w/v with 0.003% $H_2O_2$, 5-min reaction time; Sigma-Aldrich), then mounted onto gelatin-alum-coated slides and air-dried overnight. Sections were dehydrated in ascending concentration of ethanol and coverslipped with Eukitt mounting medium (Sigma-Aldrich).

#### Immunoreactivity Counting

Although a quantitative method of counting involving stereological estimates [10] is most accurate, we chose to use a semiquantitative analysis of the c-Fos immunoreactive cells involving density measurements. This method was valid because we counted the number of immunoreactive nuclei to obtain relative differences between groups and we identically sectioned brains in all experimental groups, thereby any bias in estimation in counting should be largely smaller than the biological variations[11]. An observer, blind to the experimental group assignment of each rat, quantified c-Fos immunoreactive cells under a Leica DM6000B microscope equipped with a motorized stage (x,y and z) controlled with an Antec computer with Mercator Pro software (ExploraNova). We acquired images using a Hamamatsu digital camera ORCA-03G. C-Fos neurons were considered as positive cells when the brown oval-shaped nucleus displayed staining intensity at least twice higher than the background level. For each section, we delineated the sampled area at low magnification (x2.5) and we measured the number of positive cells at high magnification (x20). The shape and the extent of the sampled area varied depending on the considered structure [9]. We then normalized the raw c-Fos positive cell counts by the sampled area (c-Fos+ cells/mm$^2$, density). To estimate the robustness of the counting method, we counted positive cells in some ROIs twice and we checked that variability between counts were inferior to 15 % with no statistical difference. We then averaged results from both counts for these ROIs.

#### Supplementary References:


1 Lenoir M, Serre F, Cantin L, Ahmed SH. Intense sweetness surpasses cocaine reward. PloS one. 2007;2(8):e698.

2 Madsen HB, Ahmed SH. Drug versus sweet reward: greater attraction to and preference for sweet versus drug cues. Addiction biology. 2015;20(3):433-44.

3 Ahmed SH, Koob GF. Transition from moderate to excessive drug intake: change in hedonic set point. Science. 1998;282(5387):298-300.

4 Ahmed SH, Kenny PJ, Koob GF, Markou A. Neurobiological evidence for hedonic allostasis associated with escalating cocaine use. Nature neuroscience. 2002;5(7):625-6.

5 Lenoir M, Guillem K, Koob GF, Ahmed SH. Drug specificity in extended access cocaine and heroin self-administration. Addiction biology. 2012;17(6):964-76.

6 Collier G, Novell K. Saccharin as a sugar surrogate. Journal of comparative and physiological psychology. 1967;64(3):401-8.

7 Smith JC, Sclafani A. Saccharin as a sugar surrogate revisited. Appetite. 2002;38(2):155-60.

8 Cantin L, Lenoir M, Augier E, Vanhille N, Dubreucq S, Serre F, et al. Cocaine is low on the value ladder of rats: possible evidence for resilience to addiction. PloS one. 2010;5(7):e11592.

9 Navailles S, Guillem K, Vouillac-Mendoza C, Ahmed SH. Coordinated Recruitment of Cortical-Subcortical Circuits and Ascending Dopamine and Serotonin Neurons During Inhibitory Control of Cocaine Seeking in Rats. Cerebral cortex. 2015;25(9):3167-81.

10 Coggeshall RE, Lekan HA. Methods for determining numbers of cells and synapses: a case for more uniform standards of review. The Journal of comparative neurology. 1996;364(1):6-15.

11 Saper CB. Any way you cut it: a new journal policy for the use of unbiased counting methods. The Journal of comparative neurology. 1996;364(1):5.

#### Supplementary Figures and Legends:

##### Figure S1

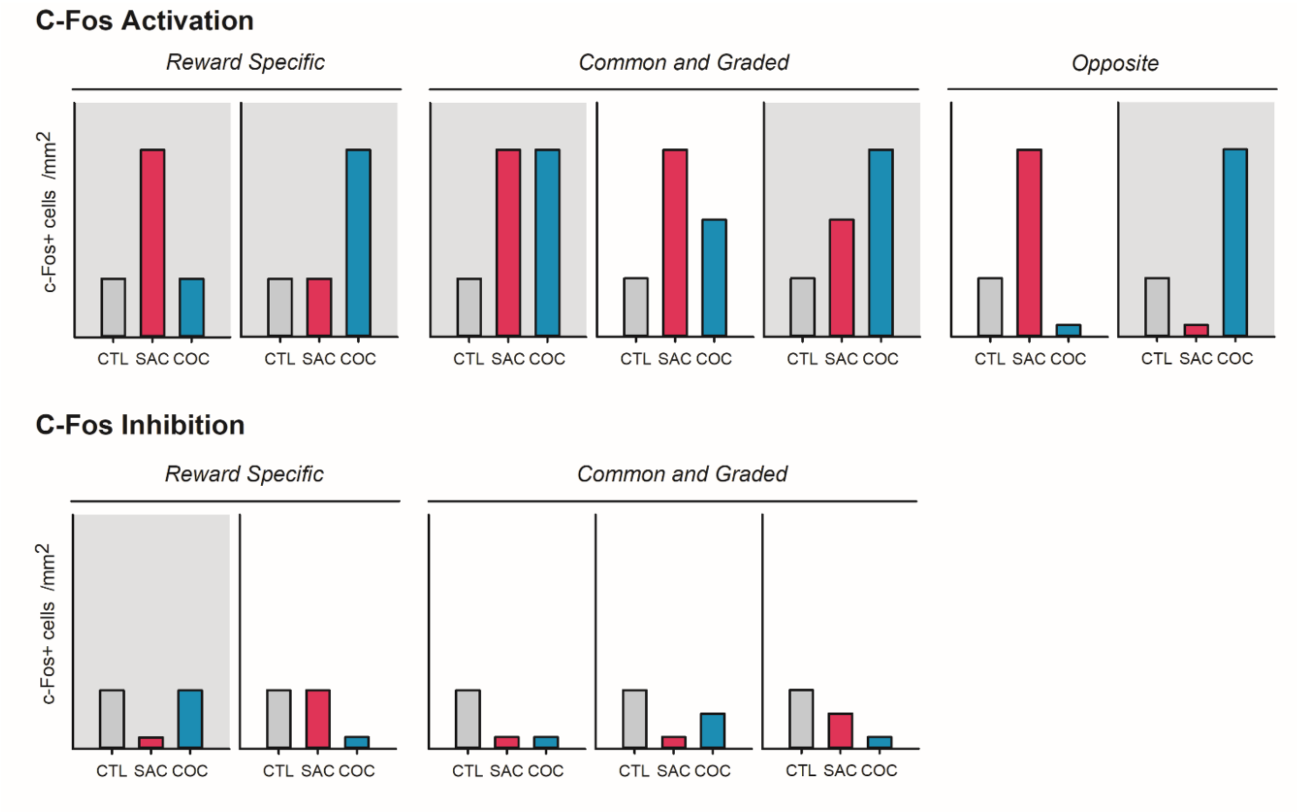


**Figure S1: C-Fos expression patterns expected in both experiments.** We expected to observe 12 potential patterns of c-Fos expression and one pattern of no group difference (not shown in this picture). Representative bar graphs show 3 main patterns: reward-specific patterns of c-Fos activation or inhibition, common and graded patterns of c-Fos activation or inhibition and opposite effects of saccharin and cocaine on c-Fos expression. Bar graphs with grey backgrounds represent patterns which were actually observed in our both experiments, whereas bar graphs with white backgrounds were not obtained. Interestingly, no pattern with significant inhibition of c-Fos in cocaine group compared to the control group was observed. CTL: control group, grey bars, rats only exposed to the operant cage (Experiment 1) or to the cue-light presentations (Experiment 2). SAC: saccharin group, red bars, rats exposed to a single trial of saccharin sampling (Experiment 1) or to a 100-min discrete sampling procedure with saccharin (Experiment 2). COC: cocaine group, blue bars, rats exposed to a single trial of cocaine sampling (Experiment 1) or to a 100-min discrete sampling procedure with cocaine (Experiment 2).

| c-Fos Patterns | ROI – Experiment 1 | ROI –Experiment 2 |
|---|---|---|
| *Reward Specific* 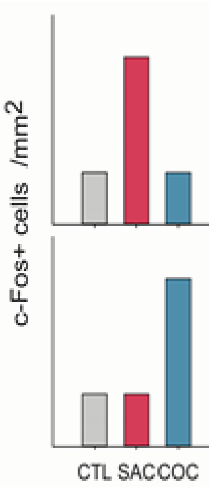  | aGC, mMS, mvDBB<br><br>rVMShell, rNAccore, mNAccore, rMS, rPaH, cPaH, **mBMA**, **mMeA**, cMeA, **mtVTA/RMTg**, **ctVTA/RMTg**, **rLDTg** | cMS, cLS, mvmVP<br><br>rVLO, mVLO, cVLO, rLO, mLO, cLO, rCg1, mCg1, mvIC, cvIC, pGC, mLShell, rDM, mDM, cDM, mDL, rDL, cDL, rVM, mVM, cVM, rVL, mVL, cVL, rCS, mCS, cCS, mdlVP, rBMA, **mBMA**, rMeA, **mMeA**, mLHb, cLHb, aSTN, pSTN, rMVTA, mMVTA, mLVTA, rtVTA/RMTg, **mtVTA/RMTg**, **ctVTA/RMTg**, mDR, **rLDTg** |
| *Common and Graded Effects* 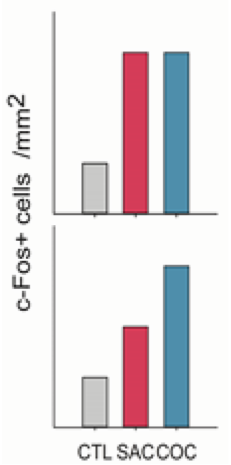  | rVO, mVLO, cCA1<br><br>rDMShell | rDMShell, VPMpc<br><br>mPPTg |
| *Opposite Effects* 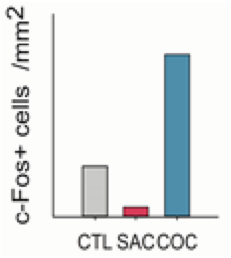  | mMVTA, rtVTA/RMTg | No Pattern |

#### Table S1:

**Patterns of cFos Activation observed in both experiments:** The table reports the subregions within the regions of interest (ROI) showing the different patterns of c-Fos activation (reward-specific, common and graded effects, and opposite effects patterns) in Experiment 1 (black color) and Experiment 2 (blue color). The subregions with a common pattern of c-Fos activation for both experiments are marked in bold. The 52 subregions within 5 major brain systems analyzed in the present study are: 1) the cortico-thalamo-hippocampal system (medial orbitofrontal cortex (MO), ventral orbitofrontal cortex (VO), ventrolateral orbitofrontal cortex (VLO), lateral orbitofrontal cortex (LO), dorsolateral orbitofrontal cortex (DLO), antero-posterior (AP) levels: +5.16, +4.68, +4.20 mm from bregma; anterior cingulate cortex (Cg1), dorsal prelimbic cortex (dPL), ventral prelimbic cortex (vPL), infralimbic cortex (IL), AP: +3.72, +3.24, +2.76 mm; ventral insular cortex (vIC), dorsal insular cortex (dIC), gustatory cortex (GC), AP: +1.44, +1.08, +0.72 mm; dorsal CA1 hippocampus (dCA1), dorsal CA3 hippocampus (dCA3), AP: -2.76, -3.24, -3.60 mm, ventral posteriomedial thalamus (VPMpc), AP: -4.08 mm), 2) the striato-pallido-septal system (dorsomedial nucleus accumbens shell (DMShell), ventromedial nucleus accumbens shell (VMShell), lateral nucleus accumbens shell (LShell), nucleus accumbens core (Core), AP: +2.76/+2.28, +1.92/+1.56, +1.20/+0.84 mm; dorsomedial striatum (DM), dorsolateral striatum (DL), ventromedial striatum (VM), ventrolateral striatum (VL), central striatum (Central), AP: +1.92/+1.56, +1.20/+0.84 mm, +0.48/+0.12 mm; medial septum (MS), lateral septum (LS), AP: +1.44, +1.08, +0.72 mm; vertical limb of the Diagonal Band of Broca (vDBB), horizontal limb of the diagonal band of Broca (HDB), AP: +1.44, +1.08, +0.72 mm; ventromedial ventral pallidum (vmVP), dorsolateral ventral pallidum (dlVP), AP: +0.36, +0.00 mm), 3) the extended and basal amygdala system (anterior bed nucleus of the striata terminalis (aBNST), AP: +0.36, +0.00, -0.36 mm, posterior bed nucleus of the striata terminalis (pBNST), ventral bed nucleus of the striata terminalis (vBNST), AP: +0.12, -0.36 mm, fusiform (Fu), AP: +0.12 mm; basolateral amygdala (BLA), basomedial amygdala (BMA), central amygdala (CeA), medial amygdala (MeA), AP: -1.92, -2.28, -2.76 mm), 4) the hypothalamic-epithalamic-subthalamic system (paraventricular hypothalamus (PaH), AP: -1.20, -1.56, -1.92 mm, arcuate hypothalamus (ArcN), AP: -1.92/-2.28, -2.64/-3.12, -3.48/-3.84 mm, ventromedial hypothalamus (VMH), AP: -1.92, -2.28/-2.64, -3.12 mm, dorsomedial hypothalamus (DMH), AP: -2.28, -2.64/-3.12, -3.48 mm; lateral hypothalamus (LH), AP: -3.12, -3.48, -3.84 mm; lateral habenula (LHb), AP: -3.12, -3.48, -3.84 mm; subthalamic nucleus (STN), AP: -3.48, -3.84 mm) and 5) the midbrain, tegmentum and pons system (medial ventral tegmental area (MVTA), lateral ventral tegmental area (LVTA), AP: -4.92, -5.28, -5.64 mm; tail of the ventral tegmental area-rostromedial tegmental nucleus (tVTA-RMTg), AP: -6.36, -6.72, -7.20 mm; dorsal raphe (DR), median raphe (MR), AP: -7.20, -7.56, -8.04 mm; pedunculopontine tegmental nucleus (PPTg), AP: -7.20, -7.56, -8.04 mm; laterodorsal tegmental nucleus (LDTg), AP: -8.40, -8.76 mm).

**Figure S2**

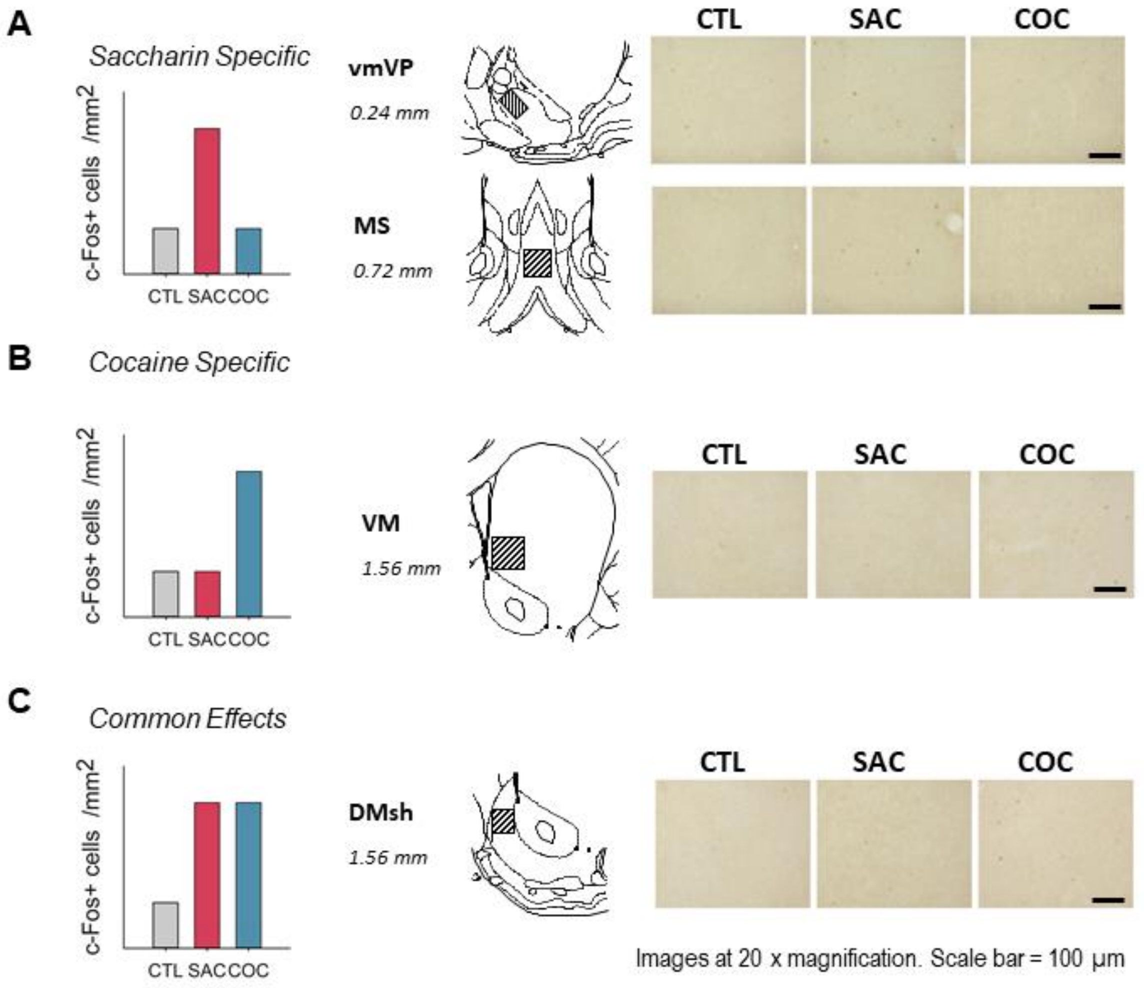


**Figure S2: Representative photomicrographs of Fos expression patterns observed in both experiments. A.** Representative bar graph shows a saccharin-specific pattern of c-Fos activation, in which c-Fos expression was significantly increased in SAC group compared with CTL and COC groups. Photomicrographs of Fos labeling in the ventromedial ventral pallidum (vmVP) and medial septum (MS) of representative rats from the CTL, SAC and COC groups showing a saccharin-specific pattern of c-Fos activation. **B.** The cocaine-specific pattern of c-Fos activation, corresponding to a significant increase in Fos expression only in the COC group compared with CTL and SAC groups, is illustrated by the representative bar graphs and the photomicrographs associated in the ventro-median striatum (VM). **C.** Example of "common effects" pattern of c-Fos activation which corresponds to a common activation of c-Fos expression in the SAC and COC groups compared with CTL group, as observed in the dorsomedial nucleus accumbens shell (DMsh). Fos-immunoreactive nuclei were counted within a squared area or the whole area, according to the brain structures, on the coronal sections taken at 0.24, 0.72 **(A)**, 1.56 **(B)** and **(C)** mm from bregma, which correspond, to the vmVP, MS, VM and DMsh (×20 magnification). Scale bar: 100 µm. See supplementary Methods for more details.

#### Figure S3

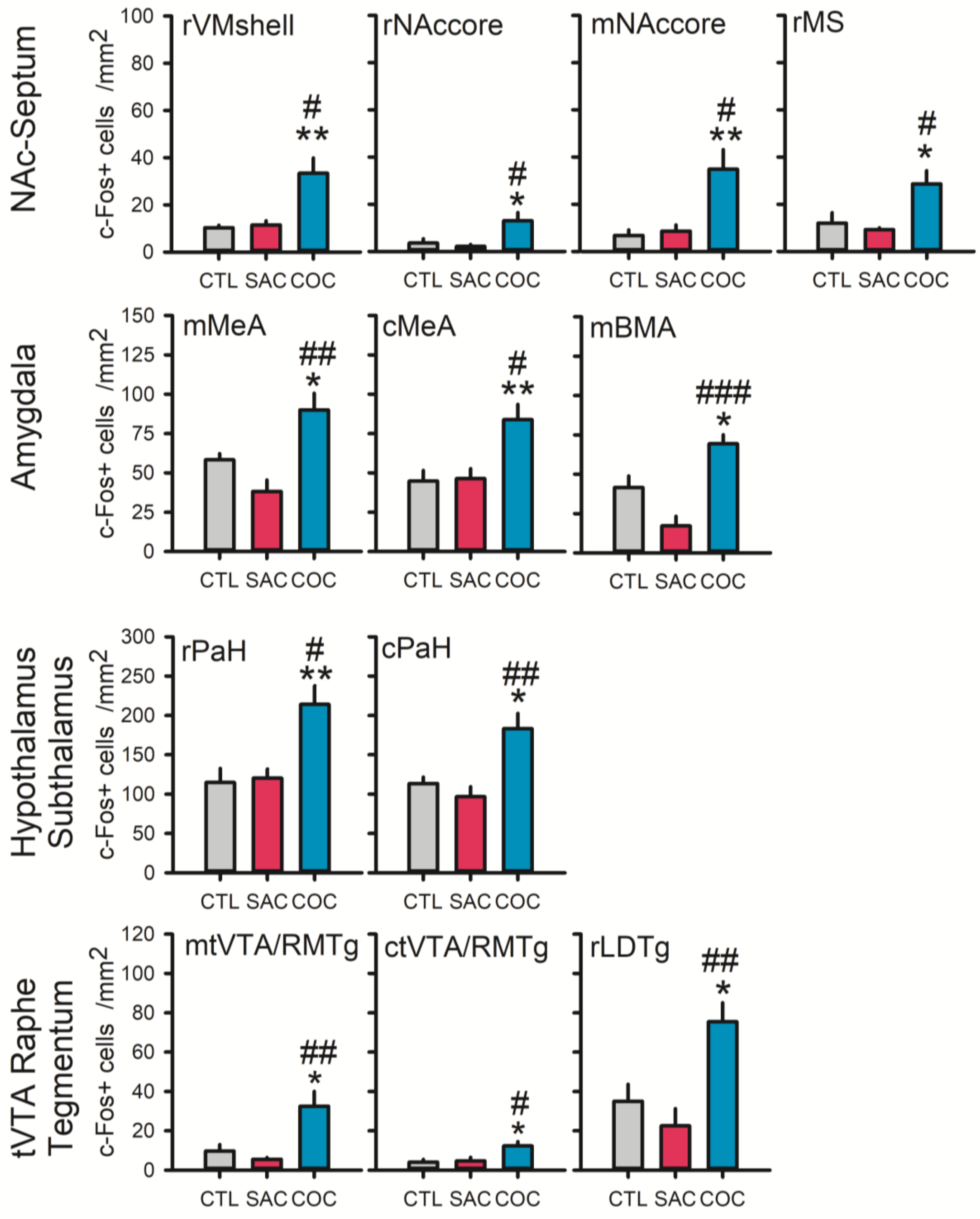


**Figure S3: Cocaine-specific patterns of c-Fos activation observed within four major brain systems in saccharin-preferring rats previously trained on the choice procedure (Experiment 1).** Bar graphs represent the mean (± SEM) density of Fos-positive cells (c-Fos+ cells /mm$^2$) counted within the brain systems (from top to bottom: striato-pallido-septal; extended and basal amygdala; hypothalamic-epithalamic-subthalamic; midbrain, tegmentum and pons systems) in rats exposed to a single trial of saccharin sampling (SAC group, red bars, n=5), a trial of cocaine sampling (COC group, blue bars, n=7) or only to the operant cage (CTL group, grey bars, n=6). * p<0.05, ** p<0.01, post-hoc

Tukey's HSD test, different from the CTL group. # $p<0.05$, ## $p<0.01$, ### $p<0.001$, post-hoc Tukey's HSD test, different from the SAC group. rVMshell, rostral level of the ventromedial nucleus accumbens shell; rNAccore, rostral part of the nucleus accumbens core; mNAccore, middle part of the nucleus accumbens core; rMS, rostral level of the medial septum; mMeA, cMeA, middle and caudal levels of the medial amygdala; mBMA, middle level of the basomedial amygdala; rPaH, cPaH, rostral and caudal levels of the paraventricular nucleus of the hypothalamus; mtVTA/RMTg, ctVTA/RMTg, middle and caudal levels of the tail of the ventral tegmental area; rLDTg, rostral level of the laterodorsal tegmental nucleus.

**Figure S4**

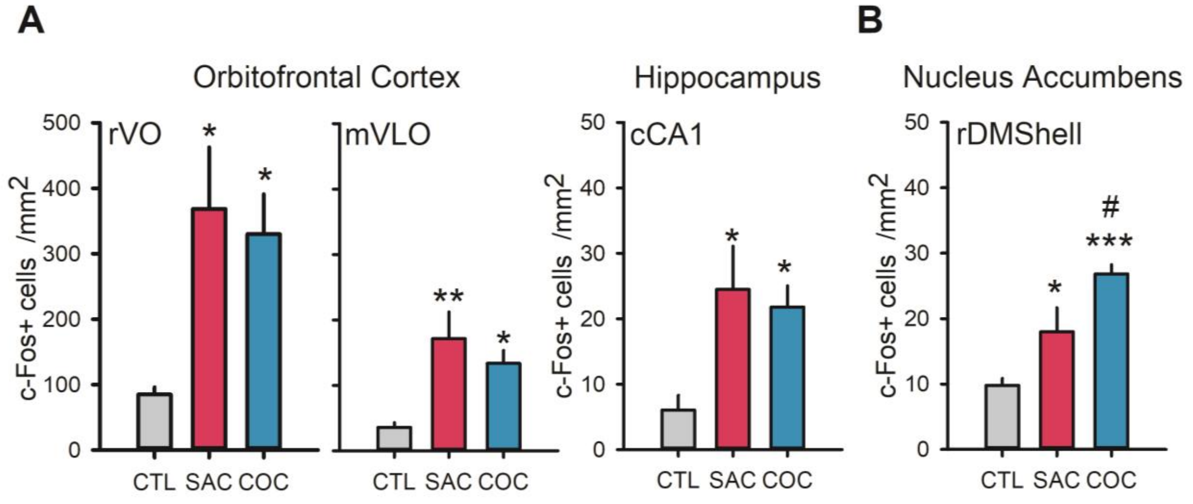


**Figure S4: Common and graded effects patterns of c-fos activation observed in brains of saccharin-preferring rats previously trained on the choice procedure (Experiment 1).** Bar graphs (mean density of Fos-positive cells ± SEM, c-Fos+ cells /mm2) show common effects **(A)** and graded effects **(B)** patterns of c-Fos activation in restricted subregions of orbitofrontal cortex, hippocampus **(A)** and nucleus accumbens **(B)** of rats exposed to a single trial of saccharin sampling (SAC group, n=5, red bars), a trial of cocaine sampling (COC group, n=7, blue bars) or only to the operant cage (CTL group, n=6, grey bars). * $p<0.05$, ** $p<0.01$, post-hoc Tukey's HSD test, different from the CTL group. # $p<0.05$, post-hoc Tukey's HSD test, different from the SAC group. rVO, rostral level of ventral orbitofrontal cortex; mVLO, middle level of ventrolateral orbitofrontal cortex; cCA1, caudal part of the dorsal hippocampal CA1 region; rDMShell, rostral level of dorsomedial nucleus accumbens shell.
